# Degradation in Binaural and Spatial Hearing, and Auditory Temporal Processing Abilities, as a Function of Aging

**DOI:** 10.1101/2024.07.08.602575

**Authors:** Carol A. Sammeth, Kerry A. Walker, Nathaniel T Greene, Achim Klug, Daniel J. Tollin

**Affiliations:** Department of Physiology & Biophysics, University of Colorado School of Medicine, Aurora, Colorado; Department of Otolaryngology – Head & Neck Surgery, University of Colorado School of Medicine, Aurora, Colorado

**Author notes:** **All correspondence should be addressed to:** Daniel J. Tollin, PhD, Department of Physiology and Biophysics, RC1-N: Rm 7106, 12800 E. 19^th^ Avenue, University of Colorado School of Medicine, Aurora, CO 80045, USA. **Human subjects IRB:** Colorado Multiple Institution IRB (COMIRB), protocol #19-1213.

**Keywords:** Presbycusis, Aging, Localization, Spatial Acuity, Spatial Hearing, Minimum Audible Angle, Spatial Hearing, Spatial Speech-in-Noise, Spatial Release from Masking, Temporal Processing, Extended High-Frequency Hearing Loss

## Abstract

**Objectives:** Sensorineural hearing loss is common with advancing age, but even with normal or near-normal hearing in older persons, performance deficits are often seen for suprathreshold listening tasks such as understanding speech in background noise or localizing sound direction. This suggests there is also a more central source of the problem. Objectives of this study were to examine as a function of age (young adult to septuagenarian) performance on: 1) a spatial acuity task examining localization ability, and a spatial speech-in-noise (SSIN) recognition task, both measured in a hemi-anechoic sound field using a circular horizontal-plane loudspeaker array, and 2) a suprathreshold auditory temporal processing task and a spectro-temporal processing task, both measured under headphones. Further, we examined any correlations between age, hearing thresholds including extended high frequency (EHF: >8000 Hz), and these measures.

**Design:** Subjects were 48 adults, aged 21 to 78, with either normal hearing or only a mild sensorineural hearing loss through 4000 Hz. The localization task measured minimum audible angle (MAA) for 500 and 4000 Hz 1/3^rd^ octave narrowband noise (NBN) in diffuse background noise for both an on-axis (reference source 0°) and off-axis (reference source 45°) listening condition at signal-to-noise ratios (SNRs) of -3, -6, -9, and -12 dB. SSIN testing was also completed for key word recognition in sentences in multi-talker babble noise; specifically, the separation between speech and noise loudspeakers was adaptively varied to determine the difference needed for 40% and 80% correct performance levels. Finally, auditory temporal processing ability was examined using the Temporal Fine Structure (TFS) test, and the Spectro-Temporal Modulation (STM) test.

**Results:** Spatial acuity was poorer (larger MAAs) in older compared to younger subjects, particularly in the more adverse listening conditions (off-axis, and poorer SNRs). The SSIN data also showed declining mean performance with age at both criterion levels, emerging in the middle age group (> 40 years), but was not correlated with standard audiometric hearing thresholds. Decreased performance on the TFS and STM tasks was dependent on age, emerging only in the older (> 60 years) and middle (>40 years) age groups, respectively; neither was dependent on hearing thresholds. Results of multiple regression analyses suggest that SSIN recognition scales with the ability of the subjects to use both low-frequency binaural temporal fine structure as well as higher-frequency binaural envelope cues, both of which are impacted by aging but not necessarily audiometric hearing thresholds. Finally, EHF range hearing thresholds significantly decreased with age, but performance on tasks remained significantly correlated with age when controlled for EHF hearing.

**Conclusions:** Particularly for more adverse listening conditions, age-related deficits, but not hearing-threshold-related deficits, were found on both of the spatial hearing tasks and in temporal and spectro-temporal processing abilities. It may be that deficits in temporal processing ability contribute to poorer spatial hearing performance in older subjects due to inaccurate coding of binaural/interaural timing information sent from the periphery to the binaural brainstem. In addition, EHF hearing loss may be a coexisting factor in the reduced performance seen in older subjects.

## INTRODUCTION

Presbycusis, or age-related hearing loss, is very common, affecting approximately one-third of adults between the age of 65 and 74, and more than half of adults aged 75 years and older (e.g., NIDCD website, 2024; ISO 2000; Cruickshanks et al. 1998). Presbycusis occurs due to degeneration of the auditory system that occurs with aging as well as the impact of a lifetime of noise exposure. It is well established that age-related hearing loss can be associated with depression, cognitive decline, dementia or Alzheimer’s disease, and generally poorer functional and psychosocial outcomes (Davidson & Guthrie 2019; Ralli et al. 2019; Rutherford et al. 2018; Zheng et al. 2017; Thomson et al. 2017; Pichora-Fuller 2015; Lin et al. 2013, 2011). Therefore, early diagnosis and treatment of hearing problems is crucial for healthy aging.

A number of mechanisms contribute to presbycusis, with the most well-known being of a peripheral nature. This includes the progressive death of outer hair cells in the cochlea resulting in hearing sensitivity loss as evidenced by elevated hearing thresholds, especially in the high frequencies (for reviews of the aging auditory system, see Fischer et al. 2020; Keithley 2020). However, even when hearing thresholds are normal or close to normal in elderly individuals, performance deficits have been reported with advanced age on suprathreshold auditory processing tasks such as understanding speech in background noise or localizing or discriminating small differences in the direction sounds are coming from, or small differences in the binaural cues to sound location (Füllgrabe et al. 2015; Freigang et al. 2014; Briley & Summerfield 2014; Dobreva et al. 2011; Dubno et al. 2008; Martin & Jerger 2005; Pichora-Fuller & Souza 2003; Abel et al. 2000; Studebaker et al. 1997; Abel & Hay 1996; Stuart & Phillips 1996), indicating that there are also other, more central underpinnings to the decreased functional auditory abilities seen with aging (see Gallun 2021 for a review). In other words, audibility and detection of sounds are not the only issue.

Central aspects of hearing deficits in the aging population are not as well understood as the peripheral and are likely multifactorial. One factor that has been implicated as potentially contributing to poorer auditory processing with aging is cognitive decline and an associated loss in working memory capacity (e.g. Lentz et al. 2022; Murphy et al. 2018; Gordon-Salant & Cole 2016; Humes et al. 2013); however, even when this is controlled, poorer performances are still seen in many older adults, suggesting it is also not the sole source of the problem. Another factor suspected to occur in the aging human auditory system, particularly in those with a significant history of noise exposure but relatively normal hearing thresholds, is cochlear synaptopathy (e.g., see Wu et al. 2019; Viana et al. 2015). Cochlear synaptopathy may occur at any age following loud noise exposures and refers to a disruption in the function of cochlear ribbon synapses between inner hair cells of the cochlea and auditory nerve fibers. This may result in subsequent neuronal degeneration and inaccurate encoding of sounds from the auditory nerve to the brainstem. Further, general neuronal degeneration with aging can occur across the auditory nervous system (Walton, 2010).

One specific hypothesis in our research laboratory’s ongoing studies on the impact of aging on hearing is that precise timing of neural activity within the binaural *brainstem* pathway declines with age (i.e., decreased synchrony). Interaural time differences (ITDs) and interaural level differences (ILDs) for sounds received binaurally vary with horizontal-plane sound source position, and these cues are encoded by the auditory brainstem circuit to assist with localization of the direction of sound as well as to perform an initial segregation of different sound sources (for reviews, see Bronkhorst 2015, 2000; Grothe et al. 2010; Colburn et al. 2006). Although there may also be age-related changes at higher levels in the ascending auditory pathway that additionally make critical contributions to source segregation (e.g. Yao et al. 2015; Middlebrooks & Bremen 2013), the brainstem is the lowest neural circuit to perform this analysis, and its outputs contribute to source segregation at the cortex (Neher et al. 2011; Grothe et al. 2010; Hawley et al. 2004). Consistent with this hypothesis is a body of evidence suggesting that auditory temporal processing ability and neural synchronization decline with age, even when the impact of peripheral hearing loss is removed or reduced, and that this decline leads to a decreased ability to segregate multiple sound sources (Füllgrabe et al. 2018, 2015; Grose et al. 2016; Ozmeral et al. 2016a,b; Gallun et al. 2014; Grose & Mamo 2012, 2010; Hopkins & Moore, 2011; Pichora-Fuller, 2003; Snell & Frisinal. 2000; Gordon-Salant & Fitzgibbons 1999; Snell 1997; Fitzgibbons & Gordon-Salant 1994). Such age-related declines reduce access to binaural cues needed for auditory space perception and good speech understanding in competing noise.

The current study examined the impact of aging on subjects ranging from young adult to septuagenarian on 2 tasks of suprathreshold spatial hearing abilities as measured in a large hemi-anechoic sound chamber, and on tests of temporal and spectro-temporal sound processing ability measured under headphones. The first spatial hearing task pertained to determining the direction from which a sound is coming. Sound localization is important for awareness of one’s environment and can be important for safety; for example, knowing the direction a vehicle is coming from when crossing a busy street. There is some evidence from previous studies of performance on sound lateralization/localization tasks that spatial acuity ability can decline in aging subjects (Srinivasan et al. 2021; Freigang et al. 2014; Briley & Summerfield 2014; Dobreva et al. 2011; Abel et al. 2000; Abel & Hay 1996). For example, examining subjects ranging in age from young adult to 81 years old, Abel et al. (2000) and Abel and Hay (1996) found poorer performance for older subjects on horizontal plane localization in sound field for broadband noise stimuli and 1/3-octave band noise centered at 500 or 4000 Hz. Dobreva et al. (2011) examined localization accuracy in the front hemifield (+90° to -90°) in adults aged 19 to 81 years old. They reported decreasing localization accuracy/precision for older adults compared to young adults for broadband, high-pass, low-pass, and narrowband noise (NBN) low-frequency targets, which the authors believed was due to age-related central declines in auditory temporal processing ability. Srinivasan et al (2021), using a headphone virtual simulation of binaural listening in the front hemifield (+90° to -90°), reported that lateralization acuity was poorer with aging for NBN at 500 and 4000 Hz.

Of note, however, is that, in these previous studies, hearing loss of up to a moderate degree was seen in some of the older subjects, which may have been a confounding factor for examining age effects. There has also been a limited number of studies on the role of aging on speech-in-noise task performance that have controlled for hearing loss, although there are a few recent studies that have done so (Patro et al., 2024; Patro et al., 2021; Srinivasan et al., 2016; Gallun et al., 2013). Therefore, in the current study, we attempted to better control the impact of peripheral hearing loss by including only subjects with either normal hearing or hearing loss of only a mild degree through 4000 Hz, so that inaudibility of the stimuli was precluded. We also excluded subjects with measurable cognitive dysfunction. Further, we used a front hemifield, horizontal spatial acuity task for both a standard on-axis listening condition (reference sound location at 0°) and a more difficult off-axis listening condition (reference sound location at 45°), with testing done in a large hemi-anechoic sound chamber. Included were both low-frequency (500 Hz) and high-frequency (4000 Hz) narrowband noise (NBN) stimuli, presented in diffuse background noise to increase task difficulty.

The second task used in the current study was a suprathreshold, Spatial Speech-In-Noise (SSIN) task. When significant peripheral hearing loss is excluded in older subjects, they are not expected to show deficits in speech recognition in *quiet* environments; however, many show deficits when listening in more *adverse* listening conditions such as competing speech or background noise (e.g., see Füllgrabe et al. 2015; Dubno et al. 2008; Martin & Jerger 2005; Pichora-Fuller & Souza 2003; Studebaker et al. 1997; Stuart & Phillips 1996). In the novel SSIN task we used, the signal-to-noise ratio (SNR) was fixed and the separation between the speech and background noise loudspeakers was varied to determine that needed for a specified percent correct performance level. A similar method was used to test SSIN performance in young normal hearing subjects by Ahrens et al (2020) and Ozmeral and Higgins (2022). It is well established that separation of an interfering noise from a target sound may result in significant spatial release from masking (SRM) in normal-hearing young listeners - - this SRM effect can be fairly well predicted from models of binaural analysis based on ITDs and ILDs (for reviews, see Colburn et al., 2006; Bronkhorst, 2000). There is some evidence that aging may result in reduced benefit from SRM (e.g. Patro et al. 2024; Gallun et al. 2013). The current study is unique in that we measured both spatial acuity ability and performance on the SSIN task in the same subjects, statistically adjusting for both age and hearing loss.

Finally, because inaccurate coding of spatial cues is expected to result in poor binaural processing ability (see Bernstein and Trahiotis, 2019), performances on 2 suprathreshold sound processing tasks were measured under headphones in this same group of subjects; specifically, the Temporal Fine Structure (TFS) test (Füllgrabe & Moore 2017) and the Spectro-Temporal Modulation (STM) test (Mehraei et al. 2014; Bernstein et al. 2013). We hypothesized that not only would poorer performance be evidenced on the suprathreshold spatial hearing tasks with advancing age, but that these might have as one underpinning a declining ability with aging for processing of temporal cues of sounds.

## MATERIALS AND METHODS

### Subjects

Subjects were 48 adults, aged 21 to 78 (35 females and 13 males). Approval was obtained from the University of Colorado Anschutz Medical Campus Institutional Review Board for use of human subjects. Subjects were paid for participation.

Inclusion criteria for subjects of all ages included either normal hearing or only a mild sensorineural hearing loss, which was defined as bilateral hearing thresholds ≤ 40 dB HL at audiometric frequencies from 250 Hz to 4000 Hz inclusively, in both ears, with no air-bone gaps > 10 dB. All subjects were also required to show relatively symmetrical hearing thresholds between the ears through 8000 Hz, which was defined as ≤ 20 dB threshold difference between the ears at all frequencies from 250 to 8000 Hz inclusively and ≤ 15 dB difference at 2 or more contiguous frequencies. All subjects had normal (Type A) tympanograms bilaterally. None of the subjects wore hearing aids.

Figure 1 shows individual audiograms and means for subjects divided into three groupings by age: younger adults (aged 20 to 39), middle-aged (aged 40 to 59), and older adults (aged 60 to 78), including hearing thresholds measured for the extended high frequency (EHF) range from 9000 to 16,000 Hz. While the older subjects do show some hearing sensitivity loss particularly in the highest audiometric frequencies, and even more in the EHF range, the audibility of the suprathreshold stimuli presented for the study was not impacted. Note that due to output limits in the EHF range for the audiometer/headphones used, a number of older subjects had no response at the maximum output level at some frequencies (maximum was 60 dB HL at 9,000 Hz, 70 dB HL at 10,000 and 11,200 Hz, 60 dB HL at 12,500. 55 dB HL at 14,000 Hz, and 30 dB HL at 16,000 Hz); When this occurred, a value 5 dB higher than that maximum was assigned to that individual for purposes of calculating the mean thresholds and for data analyses. In Figure 1, arrows indicate that a number of older subjects had no response values assigned, so that the actual mean hearing threshold at that frequency was unmeasurable but would be poorer than shown. Due to the low maximum output, 16,000 Hz is not plotted in Figure 1, but notably all younger subjects had threshold responses <30 dB HL, while some of the subjects in the middle-aged group and all of the subjects in the oldest group had no response at 30 dB HL.

**Figure 1.**
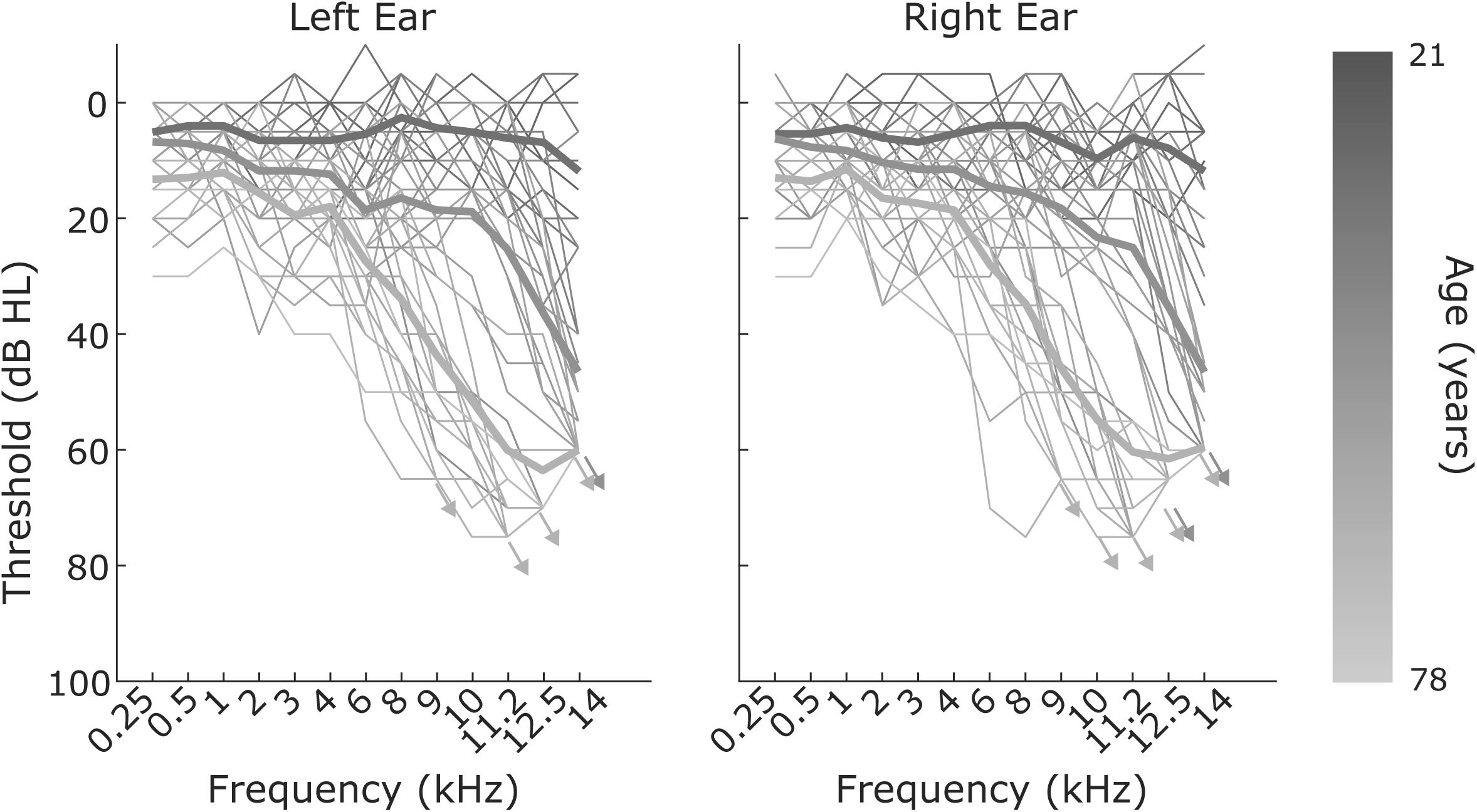
Individual audiograms and extended high-frequency hearing thresholds (in dB HL), and means for each of 3 age groupings of subjects: younger adults (aged 21-39), middle-aged (aged 42-59), and older adults (aged 60-78). For calculation of means, when a subject had no response at the maximum output limit of the audiometer at that frequency, which occurred only in the extended high frequency (EHF) range ≥9000 Hz, a value 5 dB higher was assigned (see text for more description). Arrows indicate that many older subjects had no responses at the output limits at the EHFs, so that the actual mean threshold is assumed to be poorer than that shown.

None of the subjects had a history of otologic or neurologic disorders. In addition, none of the subjects showed a cognitive deficit upon screening with the Montreal Cognitive Assessment (MOCA) test (Nasreddine et al. 2005; A passing score was taken as 26 or better correct out of 30).

Data for the spatial acuity/localization task were collected on all 48 subjects. There were more female subjects than males (35 females and 13 males); however, males were spread across both the younger and older age ranges. For a subset of 42 of the 48 subjects (31 females, 11 males), data were also available for the SSIN task. SSIN data were not available on the other 6 subjects either because their performance had previously been measured using the same test materials in a pilot study with different measurement parameters, or because they were non-native English speakers. For the Temporal Fine Structure (TFS) test, data were available for 44 of the subjects (31 females, 13 males), and for the Spectro-Temporal Modulation (STM) test, data were available for 46 of the subjects (34 females, 12 males), with the remaining subjects lost to follow-up. See Table 1 for specification of the numbers of male and female subjects per decade of life with available data for each of the measures.

**Table 1.**
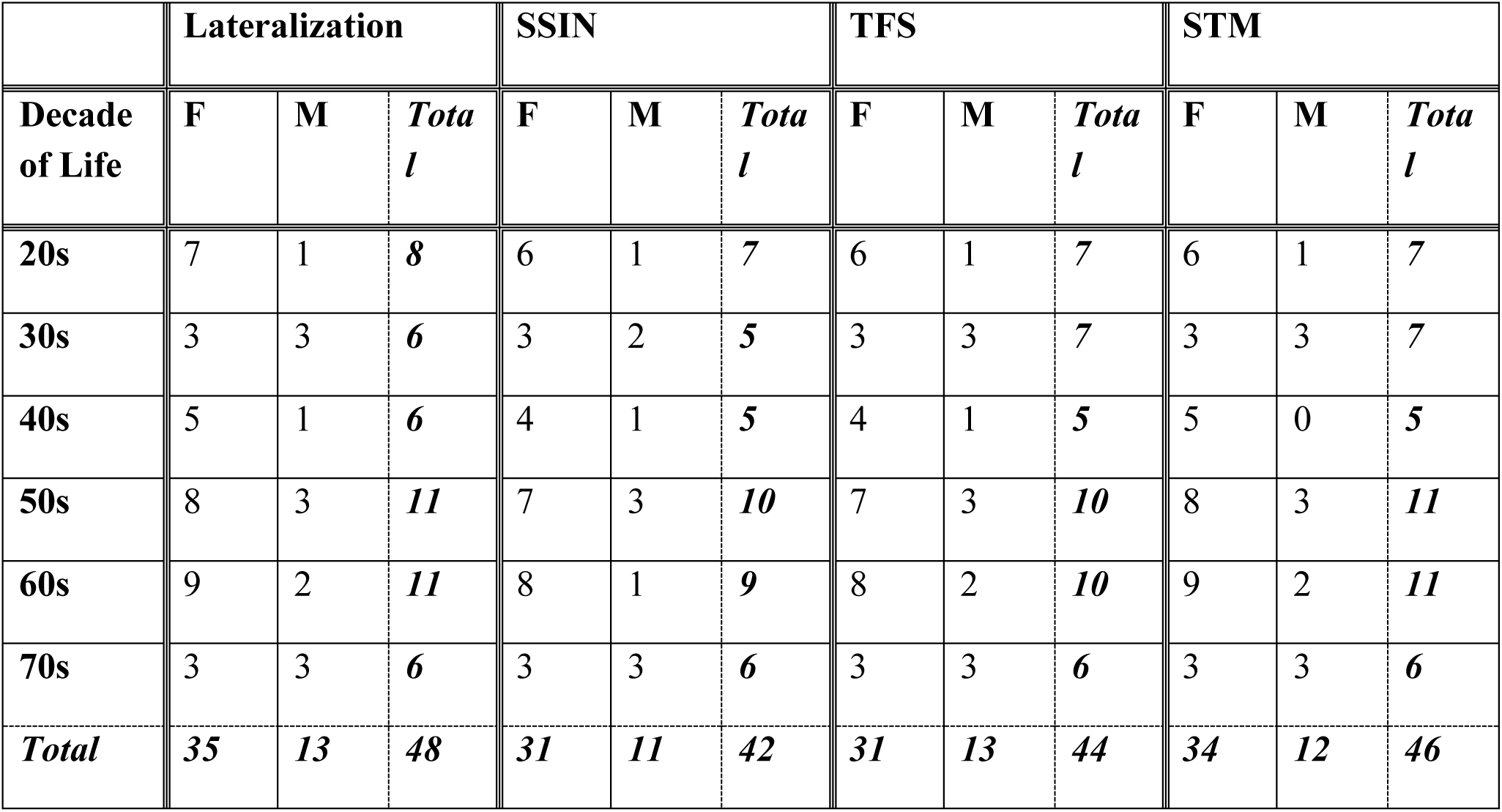
Numbers of subjects with available data for each measure by gender and decade of life. *Key:* SSIN: Spatial Speech in Noise; TFS: Temporal Fine structure; STM: Spectro-Temporal Modulation; F=female, M=male.

|  | Lateralization |  |  | SSIN |  |  | TFS |  |  | STM |  |  |
| --- | --- | --- | --- | --- | --- | --- | --- | --- | --- | --- | --- | --- |
| Decade of Life | F | M | *Total* | F | M | *Total* | F | M | *Total* | F | M | *Total* |
| 20s | 7 | 1 | 8 | 6 | 1 | 7 | 6 | 1 | 7 | 6 | 1 | 7 |
| 30s | 3 | 3 | 6 | 3 | 2 | 5 | 3 | 3 | 7 | 3 | 3 | 7 |
| 40s | 5 | 1 | 6 | 4 | 1 | 5 | 4 | 1 | 5 | 5 | 0 | 5 |
| 50s | 8 | 3 | 11 | 7 | 3 | 10 | 7 | 3 | 10 | 8 | 3 | 11 |
| 60s | 9 | 2 | 11 | 8 | 1 | 9 | 8 | 2 | 10 | 9 | 2 | 11 |
| 70s | 3 | 3 | 6 | 3 | 3 | 6 | 3 | 3 | 6 | 3 | 3 | 6 |
| *Total* | 35 | 13 | 48 | 31 | 11 | 42 | 31 | 13 | 44 | 34 | 12 | 46 |

### Equipment and Procedures

Standard air-conduction audiograms, with thresholds from 250 through 8000 Hz inclusively, were obtained with a Madsen Astera audiometer (Natus, Middleton, WI, USA) using ER-3A insert earphones (Etymotic Research, Chicago, IL, USA), as part of the enrollment screening. The EHF range was also measured from 9000 Hz through 16,000 Hz with Radioear DD450 circumaural headphones. Bone conduction thresholds were obtained at 500, 1000, 2000, and 4000 Hz via a Radioear B-71W bone oscillator. Tympanograms were obtained with an Interacoustics Titan measurement system (Middelfart, Denmark).

### Assessment of Spatial Acuity Ability

Spatial acuity ability was assessed by measuring the minimum audible angle (MAA) in the frontal horizontal plane. All testing was done in a large, hemi-anechoic sound chamber, fully treated with sound/echo-absorbing wedges except for a small part of the floor area where the subject sits, which is carpeted and has additional sound-absorbing material. A 64-loudspeaker array is available in the chamber, with stimulus presentation controlled by an Antelope Galaxy 64-channel Digital-to-Analog Convertor (Antelope Audio, San Francisco, CA, USA) and eight Crown CT8150 8-channel analog amplifiers (Crown Audio, Elkhart, IN, USA) in the control room. Loudspeakers were Gallo Acoustics Micro’s (Scotland, UK). There was a 2-way intercom between the control room and the test chamber, and a camera in the chamber for experimenter observation of the subject during testing. The camera has infrared capability for viewing the subject even when the room is darkened.

During testing, subjects were seated in a height-adjustable chair in the center of the circular horizontal loudspeaker array (Figure 2), with the chair adjusted so that the subject’s ear canal opening was at the height of the loudspeakers. Testing was done with the room dark to preclude visualization of the precise location of the loudspeakers, but an LED light came on between trials at the location of the loudspeaker directly in front of the subject, to indicate to them the orientation they should maintain during the test session. Further, the subject wore a Witmotion 901 BLE IMU position tracking headset (Guangdong, PRC) so that orientation of the forward-facing head position could be monitored during testing. The head tracker also had an LED light at the end of a frontal wand, and the subject was instructed to keep that light in line with the light on the loudspeaker in front of them throughout the testing session.

**Figure 2.**
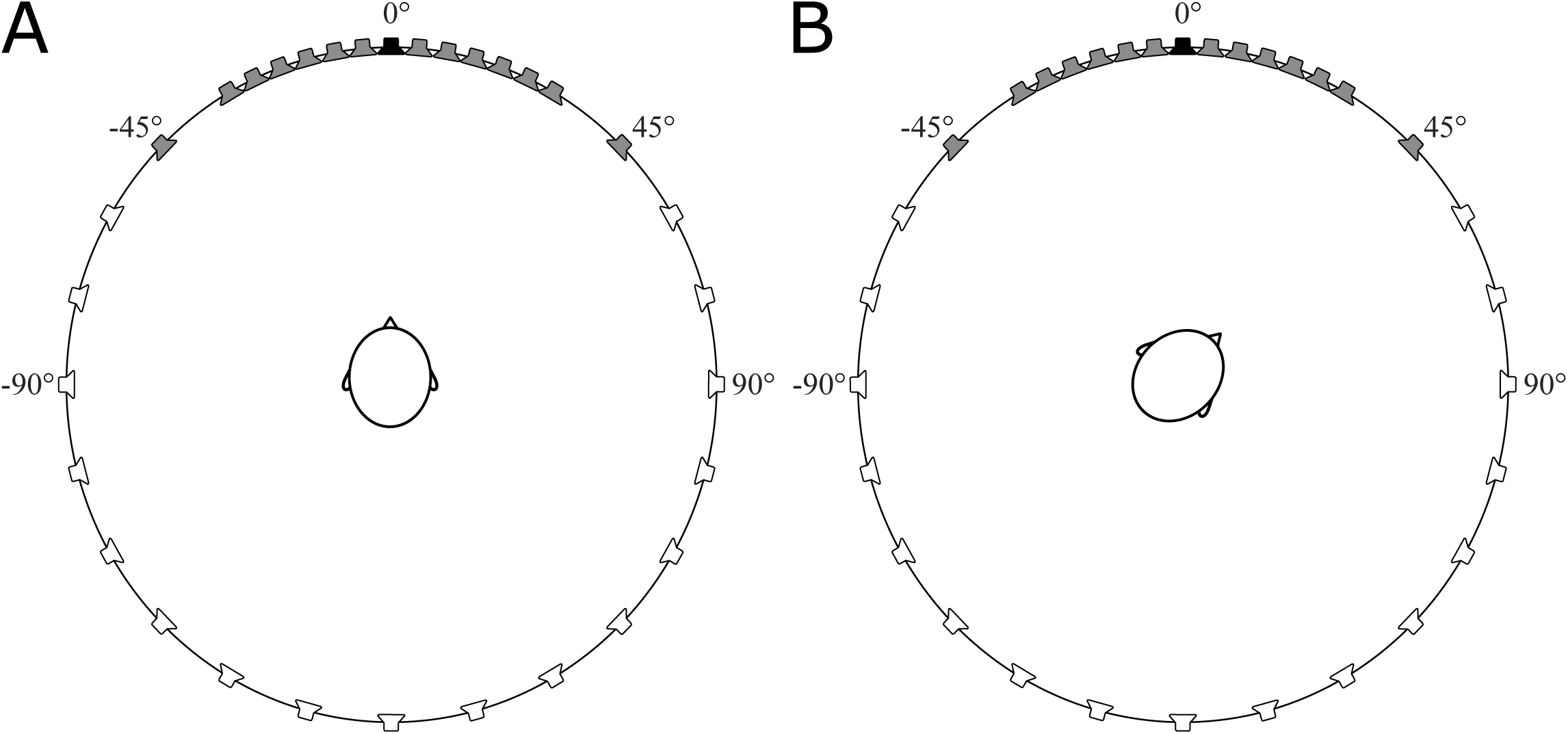
Illustration of loudspeaker spacing in the horizontal array around the subject in the hemi-anechoic test chamber. Part A: subject is seated facing 0°; dark shading indicates loudspeakers that were used for stimulus presentations for spatial acuity testing in the *on*-axis condition. Part B: subject is seated facing 45°; shading indicates loudspeakers that were used for stimulus presentations for spatial acuity testing in the *off*-axis condition.

The spatial acuity task was custom programmed using MATLAB. Stimuli were 500 and 4000 Hz narrowband noise (NBN) bursts filtered forward and backwards (using Matlab *filtfilt*) with a 1/3^rd^ octave, 4^th^ order Butterworth filter centered on the target frequency. The low-frequency and high-frequency stimuli were used to help evaluate effects of ITDs and ILDs. For pure-tone or narrowband stimuli, ITDs are known to be most effective at low frequencies (Mills, 1958, 1960). For high frequencies, the cues are ILDs and envelope ITDs. Stimuli were presented at 60 dB SPL in diffuse multi-speaker broadband noise. In separate runs, 4 fixed SNRs were tested for each stimulus: -3, -6, -9, and -12 dB; different SNRs were used in order to increase task difficulty and to find a test that was useful in separating out performance as a function of age. The order of testing of the 2 stimulus frequencies was randomized across subjects and sessions, with SNR progressively worsening (from -3 to -12 dB) with each run. Practice runs were used to ensure that the subject understood the task and was familiar with the sound of each NBN stimuli before beginning test runs.

Testing was completed across two 2-hour sessions. In the first session, all testing was done in the “on-axis” listening condition; that is, the subject was seated facing the loudspeaker at 0°. The set-up is illustrated in Figure 2, part A - - used for target stimuli for this task were the darker shaded loudspeakers from +45 to -45° inclusively. Loudspeakers were spaced 5° apart from -30° to +30°, which covered the range of final MAA values for the vast majority of subjects, with slightly larger spacing for the 2 wider angle loudspeakers. Noise emanated from loudspeakers placed at 30° to 330°, in 30° increments, but was never co-located with the target stimuli.

A 2-alternative forced choice (2AFC) procedure was used. Following the classic MAA task paradigm of Mills (1958), the first stimulus was always presented from the loudspeaker at 0° (directly in front of the subject) and the second stimulus was presented from a loudspeaker within +/- 45°. The subject’s task, using 2 buttons on a handheld keypad, was simply to indicate if they believed the second stimulus had been presented to the right or to the left of the first stimulus. A 2-down, 1-up procedure (Levitt, 1971) was used for each run (2 correct responses before a decrease in step size, but an increase in step size for every incorrect response) to determine the 70.7% correct point. Two runs were interleaved (i.e., odd trials corresponded to track 1 and even trials corresponded to track 2) to avoid any pattern being perceived by the subject. The first presentation of the 2^nd^ stimulus was always from a loudspeaker at 45° (+/-), an easy task, then presentations continued to smaller angles, with an initial 2-speaker step size for 3 reversals (change in direction of the stimulus angle difference), and then a single speaker step size for 6 more reversals. Whether the 2^nd^ stimulus was presented to the right or left of the first (+/-) was randomized. Note that, if the subject’s performance was excellent, there was the possibility of no angle difference (i.e., both presentations at 0°), which always counted as an incorrect response and a reversal (this occurred most often for younger subjects and easier task conditions).

The MAAs for each of the 2 interleaved runs, calculated as the mean of the last six reversals, were averaged to obtain the final MAA value that was used in analyses of the data. If the 2 interleaved runs showed very disparate values (which happened occasionally when a subject became tired and their attention momentarily wandered), another set was done after the subject took a rest break, and the 2 lowest MAA values of the 4 were the ones averaged for analysis. Finally, following Hartmann & Rakerd (1989), who demonstrated that the Mills (1958) method to measure MAA overestimates subject performance, to compute our final estimate of MMA we multiplied the estimate computed above by a factor of 2.

In a second session done on a different day, the same testing was done but in the more difficult, “off-axis” listening condition, i.e., the subject was seated facing the 45° loudspeaker as illustrated in Figure 2, Part B. The first stimulus was still presented from the loudspeaker at 0° (in this off-axis condition, this is at -45° relative to the subject’s gaze) and the second stimulus was still presented from a loudspeaker within +/- 45° of the initial presentation (that is, from 0° to -90° relative to the subject’s gaze). Thus, the stimuli for the off-axis condition were presented relative to the reference speaker at 45°, toward the left ear of the subject.

### Assessment of Spatial Speech-In-Noise (SSIN) ability

The SSIN task was completed in the hemi-anechoic sound chamber in another session, on a different day, with the lights in the test chamber remaining on during testing. For this testing, subjects were also seated in the chair in the center of the circular horizontal loudspeaker array with loudspeakers at ear height. The SSIN task was custom programmed using MATLAB. Assessed was the ability of the subject to repeat target words from the Harvard IEEE (IEEE 1969) corpus of sentences. These stimuli represent everyday conversational but low-contextual speech, and each sentence has 5 key words. The sentence materials and their recording method are described in Nelson et al. (2003); however, for the current study, only the sentences spoken by a *female* native American English speaker were used. The background noise we used was six-talker babble from the Connected Speech Test (Cox et al., 1987). The audio file is publicly available at the HARL Memphis website - https://harlmemphis.org/connected-speech-test-cst/. Calibration was done by generating a noise stimulus with the same spectrum as the long-term average speech spectrum of the experimental stimuli (IEEE sentences, or babble noise) and measuring the sound pressure level of the noise in dB(A). For the current study, the IEEE sentences were presented at an average conversational level of 60 dB(A).

Four fixed SNR conditions were tested in separate runs: -3, -6, -9, and -12 dB. Additionally, 2 tracks were interleaved within a single test condition: one track was based on a 40% correct performance criterion (≥2 of the 5 key words repeated correctly), and the second track was based on an 80% correct performance criterion (≥4 of the 5 key words repeated correctly). For each presentation, subjects were instructed to repeat aloud as many of the words in the sentence as possible, and guessing was encouraged.

The sentences were always presented from 0° azimuth (the loudspeaker directly in front of the subject), and the babble noise was presented from another loudspeaker located within +/- 90° of the subject on the horizontal axis with random positive or negative presentations. Available *noise* loudspeakers for this task were at the following azimuths: 0°, +/-5°, +/-10°, +/-15°, +/-20°, +/-25°, +/-30°, +/-45°, +/-60°, +/-75°, and +/-90°. The separation between the signal and noise loudspeaker was modified adaptively, starting at +/-90° and proceeding down to as small as a 5° difference, to assess the subject’s ability to accurately separate the target speech from the background babble. Specifically, the program first decreased or increased the spatial separation (based on the track criterion) by 3 loudspeakers until 4 reversals were observed in a track, then the program moved to a single speaker step size. Testing continued for 20 sentences per track (40 sentences total). The spatial separation of the speech and noise loudspeakers that was needed to achieve 40% and 80% correct for each SNR condition was calculated by averaging across the final 10 trial locations for each track.

It should be noted here that this task presents the maskers from only one side of the subject, with side chosen randomly from trial to trial, with the target being always presented from the midline. As such, the necessity of subjects to use binaural cues for the task might be reduced because the subjects potentially have access to monaural better ear glimpsing of the speech, that is, attending to the ear with the better SNR (Brungart and Iyer, 2012). The better ear advantage is particularly strong when the target and masker spatial separations are large, such as would be the case with our midline target and a masker at 90° (Bronkhorst & Plomp, 1988). In this case, the head shadow effect creates an SNR advantage at the ear opposite the masker, and thus information may be available that does not rely on binaural processing per se. However, the better ear effect, while largest for large target to masker spatial separations, becomes less effective for separations < ∼45° (Bronkhorst & Plomp, 1988) as the head shadow magnitude becomes much smaller. In these conditions, subjects potentially rely more on binaural cues. Out of 168 total SSIN measurements in all subjects with all four SNRs in the 80% correct case (see Figure 5B), 143 measurements (85%) had spatial separation thresholds of <45°, and 77% were < 30°. Thus, it is likely that for the bulk of our measurements in this SSIN task, subjects were utilizing binaural cues, and occasionally better ear glimpsing as well.

### Assessment of Temporal and Spectro-temporal Sound Processing Ability

Two suprathreshold psychoacoustic tests were completed with testing in a sound-treated booth under Sennheiser HD-25 headphones (Wedemark, Germany). These tasks were custom programmed using MATLAB software (Natick, MA, USA), with stimuli delivered binaurally from a desktop PC in the control room via use of a Fireface UCX Digital-to-Analog Convertor (RME, Haimhausen, Germany) and Magni 3 amplifiers (Schitt Audio, Newhall, CA, USA) to the headphones. Subjects sat at a desk and used a computer mouse to indicate their responses on a monitor screen placed in front of them. The installation files for these tests were downloaded from the University of Cambridge Auditory Perception Group in the Department of Psychology webpage (https://www.psychol.cam.ac.uk/hearing, 2018).

The first task was the Temporal Fine Structure - Adaptive Frequency (TFS-AF) test. The stimuli and procedures used were identical to those of Füllgrabe and Moore (2017). The TFS-AF assessment uses an adaptive, two-interval, 2AFC procedure, with visual feedback for listener responses. The objective is to determine the highest frequency at which a listener can perceive a fixed interaural phase difference (IPD). Thus, a higher value indicates better performance.

Specifically, for this task, the stimulus frequency was adaptively modified using a 2-down, 1-up procedure to determine 70.7% correct performance (Levitt, 1971). Following Füllgrabe and Moore (2017), the starting frequency was 200 Hz, with frequency increasing by a factor of 1.4 until the first reversal, 1.2 until the second reversal and by a factor of 1.1 for the remainder of the reversals. Stimuli were presented at each test frequency at approximately 30 dB SL (Sensation Level, i.e., dB above each individual’s measured hearing thresholds), and 2 intervals were presented sequentially with a 500 msec gap between each presentation. Intervals consisted of four, 400-msec pure tones presented successively and separated by 100 msec gaps. For each trial, a target and standard interval were randomly generated. The standard interval consisted of 4 pure tones at a given frequency with a 0-degree IPD, which resulted in a stationary perception in the head (diotic presentation). In contrast, the target interval added a fixed IPD for the second and fourth pure tones to create the perception of a diffuse, moving pure tone within the listener’s head (dichotic presentation). In separate blocks, 3 different IPDs were tested in each subject, 30, 60 and 180°. On the computer monitor, the listener was presented with 2 boxes, labeled ‘1’ and ‘2’, which were highlighted during their corresponding stimulus presentation interval. Listeners were instructed to select the interval that corresponded to the perception of a sound that was moving. Feedback was given as a green box containing the word “correct” or a red box containing the word “incorrect”. Following eight reversals, the trial ceased and a geometric mean of the final six reversal frequencies resulted in a threshold estimate for the given IPD.

The second task used was the Spectro-Temporal Modulation (STM) test (for a detailed description, see Bernstein et al. 2013). This task is also an adaptive 2AFC assessment. It uses diotic presentation as it applies a set temporal modulation and spectral density to an octave-band noise carrier, which is the same at both ears. The STM evaluates the minimum modulation depth the listener can detect, and thus, for this task, a lower or more negative number indicates better performance. Stimuli were presented at 85 dB(A) over the headphones at a set four-octave noise carrier, spectral ripple density (cycles per octave; c/o), modulation rate (in Hz), and direction. There were 3 different modulation conditions in the testing protocol: 1) 1000 Hz center frequency, 2 c/o, 4 Hz, upward direction, 2) 4000 Hz center frequency, 4 c/o, 4 Hz upward direction, and 3) broadband noise, 2 c/o, 4 Hz upward direction.

In this task, listeners were presented with a pair of sequential stimuli, one consisting of noise and the other containing the set modulation. Stimuli were 500ms in duration and presentations were separated by 200 ms silence gaps. The starting modulation depth was 0 dB and varied by a three-down, one-up adaptive track that resulted in a 79.4% correct point (Levitt, 1971). Feedback was provided to the listener visually with “correct” and “incorrect” text prompts. Trials consisted of 9 reversals overall, whereas depth was initially adjusted by 6 dB then 4 dB following the first reversal, and then by 2 dB following the third reversal. The average modulation depth across the last six 2-dB step size reversals was used to calculate the modulation threshold.

### Statistical Analyses

Analyses of the data included primarily age as a continuous variable. Correlation analyses using the Pearson product-moment correlation coefficient (r) were used. Both multiple linear regression and partial correlation analyses were performed for a number of different predicted variables and a number of different predictors. Because a number of the predictors were often highly correlated, such as age and hearing loss, the results of partial correlation analyses are mainly presented here. However, for most analyses, both partial correlation and multiple regression analyses showed the same trends. Significance of tests were taken as p < 0.05. To examine the effects of age group and task difficulty conditions (e.g., different SNRs) on performance, one-way or two-way analyses of variance (ANOVAs) were performed with Bonferroni’s multiple comparisons test serving as the post-hoc analysis method. For graphical purposes such as examination by age groupings, results are displayed as box-and-whisker plots, which show the median, lower, and upper quartiles, and variability of the data. All statistical tests were performed using Graphpad Prism v.10 statistical software.

## RESULTS

### Audiometric Thresholds

Hearing thresholds were significantly correlated with age for all frequencies from 500 Hz to 14 kHz inclusively: r values ranged from 0.53-0.88 (p < 0.0001) with r increasing systematically with frequency. As grouped, the audiometric 4-frequency pure tone average (PTA; 500, 1000, 2000 and 4000 Hz) (r = 0.68, p < 0.0001) and averaged EHF thresholds (EHFA; 9000-14000 Hz) (r = 0.87, p < 0.0001) were also significantly correlated with age. Across the population, PTA had a mean of 9.83, median of 8.33 and a range of 0 to 30 dB HL; EHFA had a mean of 32.35, median of 29.75 and range of -0.5 to 68 dB HL. Forty-five of the 48 subjects had PTA ≤ 20 dB HL.

To examine at what age PTA thresholds begin to increase, a one-way ANOVA revealed a significant effect of age group [F(2,45) = 13.34, p < 0.0001] with post-hoc tests showing significant differences between the older age group and both the young (p < 0.0001) and middle (p = 0.01) age groups, but no difference between the young and middle ages (p = 0.08). Similar analysis was performed with EHFA thresholds (8000-14,000 Hz), with post-hoc tests revealing significant pairwise differences between all 3 age groups (p < 0.0001). Thus, in this cohort of subjects, decreases in hearing thresholds in the standard audiometric frequency range (PTA) begin to emerge only in the older age group (> 60 years), but poorer thresholds in the EHFA range begin to emerge already in the middle age group (> 40 years).

Finally, as expected from the results above, EHF and PTA thresholds were also highly correlated (r = 0.72, p < 0.0001). To control for the strong age effects on EHF and PTA thresholds, partial correlation analyses were performed. Results reveal that the correlation between EHF and PTA thresholds was still significant even when controlling for age (r = 0.36, p = 0.013). This suggests that the EHF hearing loss exhibited already in the middle age group is partly predictable by the PTA thresholds.

### Spatial Acuity as a Function of Age

Shown in the 2 left panels of Figure 3 are scatterplots of individual MAA results as a function of age for each stimulus (Part A: 500 Hz and Part B: 4000 Hz) at each of the 4 SNRs for the *on*-axis (0°) subject orientation. A larger value indicates poorer performance on this task. Solid lines indicate the linear regression trends. The right panels in Figure 3 are box-and-whisker plots of MAA results at each SNR for the 3 age groupings: younger adults (n = 14), the middle group (n = 17) and the older adults (n = 17). As can be seen, there was a slight reduction in performance for a few subjects older than age 60. MAA with 500 Hz was significantly correlated with age for all SNRs (r’s 0.3-0.49, p < 0.05). Greater performance variability was found across all ages for the 4000 Hz stimuli (bottom row). MAAs for 4000 Hz were not correlated with age for any SNR. Most subjects had an average calculated MAA <10° for 500 Hz and <20° for 4000 Hz, indicating this was a relatively easy task for either stimulus.

**Figure 3.**
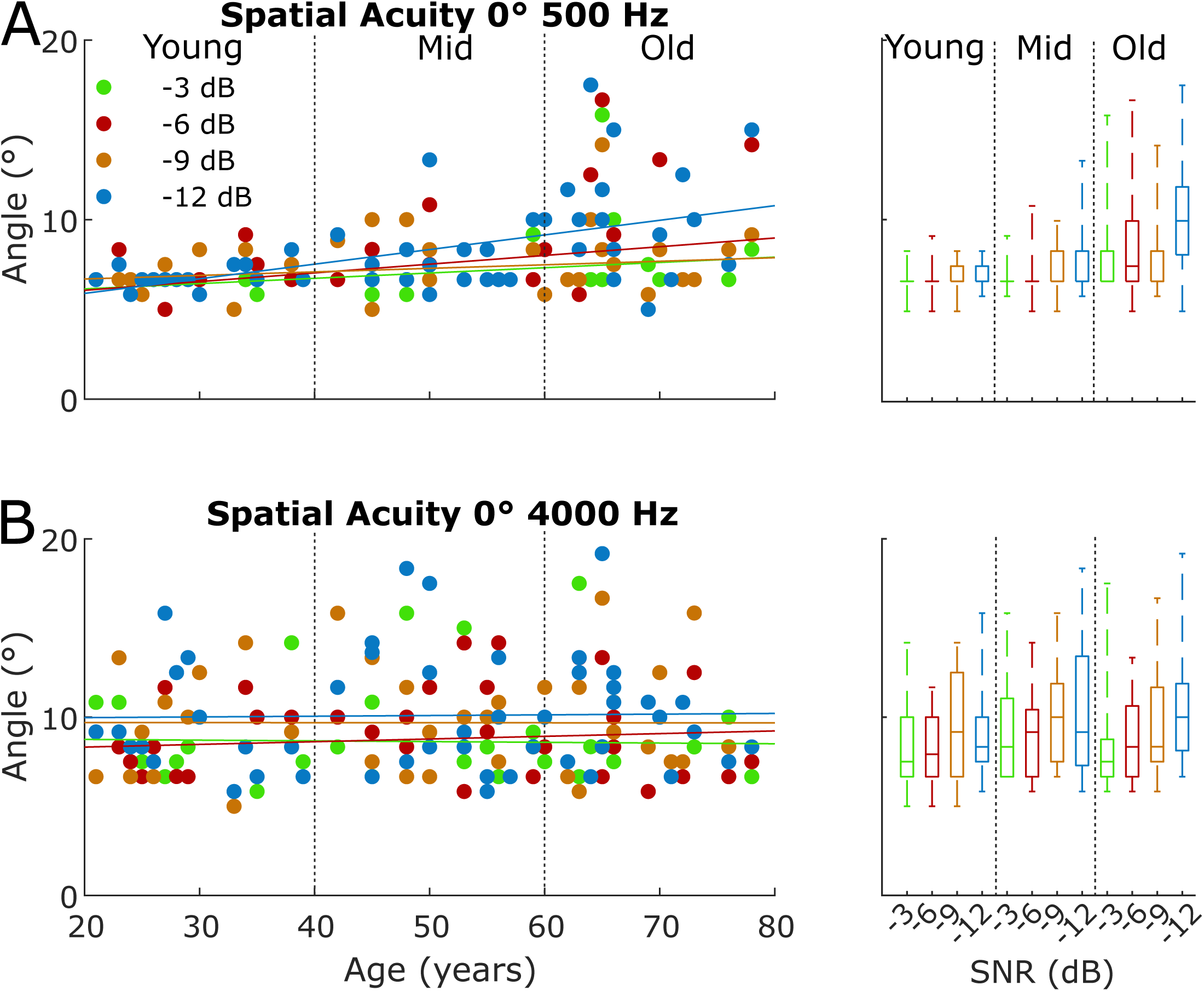
Spatial acuity/localization data for the *on*-axis listening condition (subject seated facing 0°) for each dB SNR (-3, -6, -9, -12). Part A (top row) shows results for the 500 Hz stimulus and Part B (bottom row) for the 4000 Hz stimulus. Left panels show individual data scatterplots of minimum audible angle (MAA, in degrees) by age (in years). A lower MAA value indicates better performance. Solid lines indicate the linear regression trend for each SNR. Right panels show box-and-whisker plots with the median, lower and upper quartiles, and variability of the data, for 3 age groupings: younger adults (aged 21-39), middle-aged (42-59), and older adults (60-78).

To examine the contributions of age and 500 Hz audiometric thresholds to 500 Hz MAAs, partial correlation analyses were performed. Both MAA versus age and versus threshold were controlled for by threshold and age, respectively. Partial correlations were used because audiometric thresholds were significantly correlated with age for all frequencies, making it difficult to interpret the results of multiple regression analyses. The only condition that survived this analysis was for -12 dB SNR where there was a significant correlation between MAA and age (r = 0.42, p = 0.003). For the 4000 Hz stimulus, there were no significant correlations between age and MAA at any SNR.

Figure 4 shows comparable data but for the harder condition of the *off*-axis, or within hemifield, subject orientation. *Note that the vertical y-axis ranges are different for on-axis condition plots (Figure 3) versus the off-axis condition plots (Figure 4) in order to best display the data*. The scatterplots for this harder, off-axis condition show poorer performances in general than for the on-axis orientation for both the 500 Hz stimulus (Part A) and the 4000 Hz stimulus (Part B).

**Figure 4.**
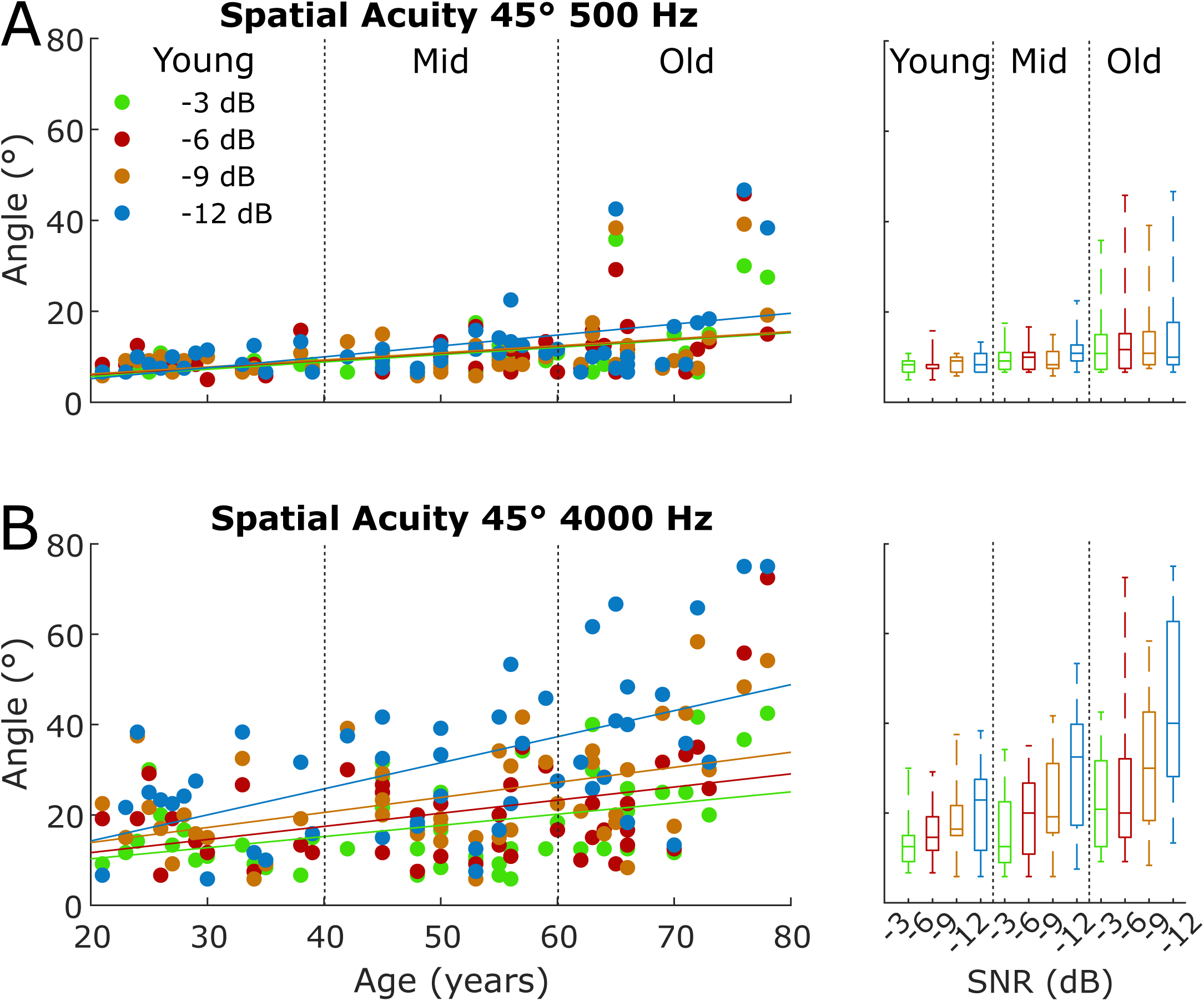
Spatial acuity/localization data for the *off*-axis listening condition for each dB SNR (-(- 3, -6, -9, -12). Note the vertical axis change relative to Figure 3. Part A shows results for the 500 Hz stimulus and Part B for the 4000 Hz stimulus. Left panels show individual data scatterplots of minimum audible angle (MAA, in degrees) by age (in years). A lower MAA value indicates better performance. Solid lines indicate the linear regression trend for each SNR. Right panels show box-and-whisker plots with the median, lower and upper quartiles, and variability of the data, for 3 age groupings: younger adults (aged 21-39), middle-aged (42-59), and older adults (60-78).

Further, the impact of aging is more pronounced in this condition, especially at the 2 poorest SNRs (-9 and -12 dB) for 4000 Hz stimuli, with the 2 oldest subjects showing large MAA values ranging from 46° to 74°. Even young subjects had more difficulty in the off-axis condition for the 4000 Hz stimulus than for the 500 Hz stimulus, but variability in performances was notably higher for older subjects.

A two-way ANOVA was conducted that examined the effect of age group and SNR on MAA with 500 Hz. There was no significant interaction between the effects of age group and SNR on MAA [F(6, 180) = 0.09, p = 0.99]. Simple main effects analysis showed significant effects of age (p < 0.0001) but not SNR (p =0.51). These results suggest that spatial acuity for off-axis 500 Hz worsens with age but not SNR. The same analysis was performed for the 4000 Hz condition. Here there was also no significant interaction between the effects of age group and SNR on MAA [F(6, 180) = 0.89, p = 0.51]. Simple main effects analysis showed significant effects of both age (p < 0.0001) and SNR (p < 0.0001). These results indicate that spatial acuity for off-axis 4000 Hz declined comparably with SNR across age group.

To investigate the factors involved in the above results, additional analyses were conducted. Consistent with the above results, off-axis MAAs for both 500 Hz and 4000 Hz and at all SNRs were significantly correlated with age (r values ranged from 0.38-0.54, p < 0.007). Similar to the above, partial correlation analyses were performed to disentangle effects of age and threshold. For the 500 Hz off-axis MAA, only the -6 dB SNR condition was significant between MAA and age controlling for 500 Hz hearing threshold (r = 0.31, p = 0.034). None of the correlations were significant when comparing MAA and threshold controlling for age. For the 4000 Hz stimulus, only the -12 dB SNR condition was significant between MAA and threshold at 4000 Hz (r = 0.5, p = 0.0003) when controlling for age.

To examine the age at which performance declines began to emerge, a one-way ANOVA for the 500 Hz -12 dB SNR midline MAA condition resulted in a significant difference in MAA across the 3 age groups [F(2,45) = 9.8, p = 0.0003]. Post-hoc tests revealed significant pairwise differences between the older subjects and both the young (p = 0.0004) and middle (p = 0.006) age subjects but no significant difference between young and middle aged (p = 0.52). The same conclusion was found in analyses of the 4000 Hz -12 dB SNR off-axis condition indicating that declines in spatial acuity at 500 Hz and 4000 Hz, and thus the capability to utilize low-frequency fine structure ITD and high-frequency envelope ITDs, respectively, only begin to emerge in the older subjects > 60 years.

### Spatial Speech-in-Noise by Age

Shown in the left panels of Figure 5 are scatterplots of individual SSIN results (i.e., speaker separation needed, in degrees, for the given percent correct) as a function of age for each of the 4 SNRs for 40% correct performance (Part A) and for the harder 80% correct performance criterion (Part B). A larger value indicates poorer performance on this task. Solid lines indicate the linear regression trends. The right panels in Figure 5 are box-and-whisker plots of SSIN results for the 3 age groupings: younger adults (n = 12), the middle group (n = 15) and the older adults (n = 15). It can be seen that performance generally decreased with decreasing SNR. Performance also tended to decline with advancing age for both percent correct performance criteria, with, again, greater variability in performance for older than for younger subjects. Performance on the SSIN task was significantly correlated with age for both the 40% and 80% correct criterion and all SNRs (r’s ranging 0.3-0.55, p < 0.05).

**Figure 5.**
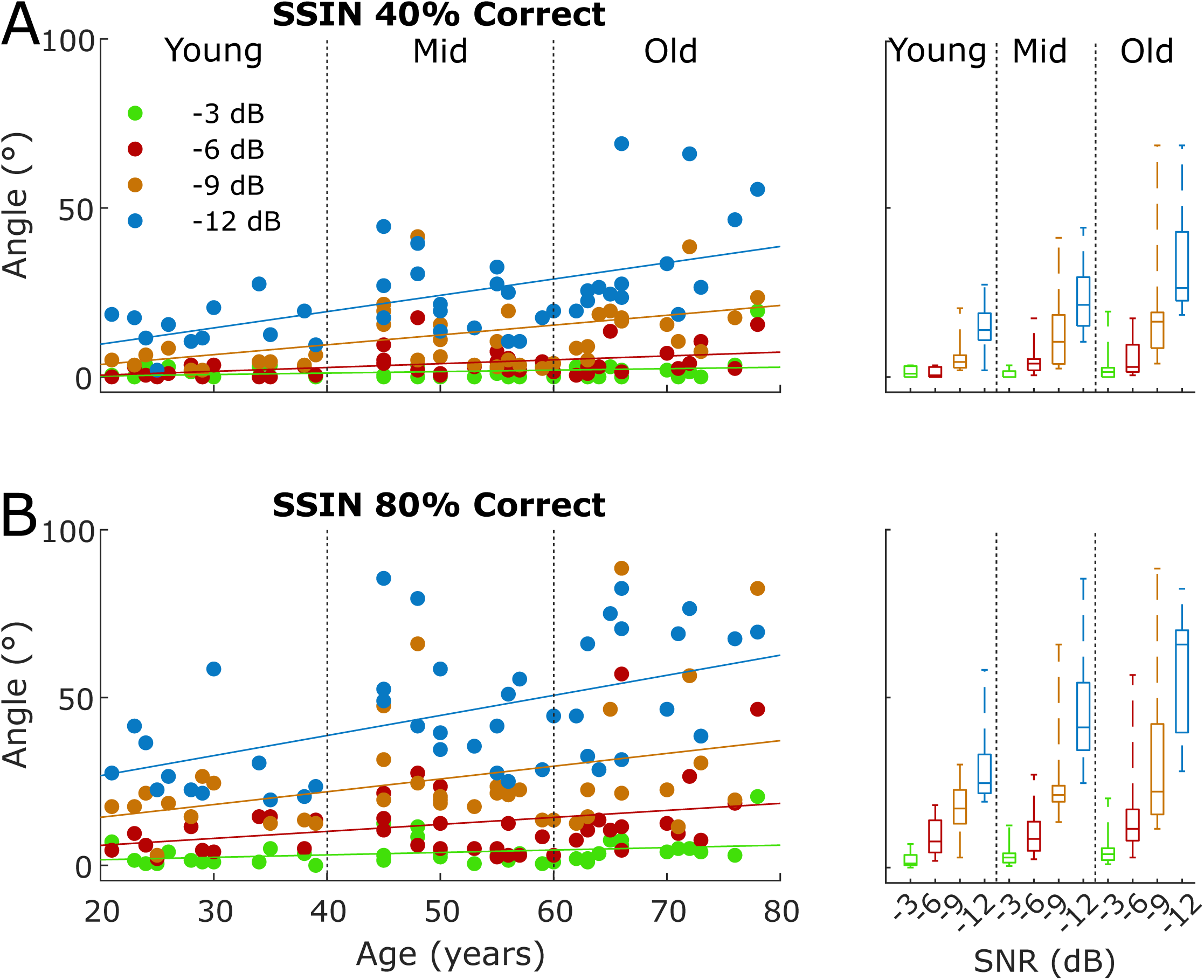
Spatial speech-in-noise (SSIN) data. Part A shows results for the 40% correct criterion and Part B for the 80% correct criterion, for each dB SNR (-3, -6, -9, -12). Left panels show individual data scatterplots of the speech and noise loudspeaker separation needed (angle, in degrees) for the given performance criterion by age (in years). A lower value indicates better performance. Solid lines indicate the linear regression trend for each SNR. Right panels show box-and-whisker plots with the median, lower and upper quartiles, and variability of the data, for 3 age groupings: younger adults (aged 21-39), middle-aged (42-59), and older adults (60-78).

A two-way ANOVA was conducted that examined the effect of age group and SNR on performance in the SSIN task at the 80% correct criterion. There was a slight, but statistically significant, interaction between the effects of age group and SNR on threshold [F(6, 156) = 2.2, p = 0.045]. Simple main effects analysis showed significant effects of both age (p < 0.0001) and SNR (p < 0.0001). These results suggest that performance in the SSIN task worsens with SNR and also age, but that the decline in performance is larger in older subjects than in younger. Similar results were found for the 40% correct SSIN task (not reported).

Additional analyses were performed to investigate the factors that might be involved in the performance of the SSIN task. For the 40%-correct criterion, partial correlation analyses controlling respectively for age or PTA threshold revealed a significant correlation between SSIN at -6 dB SNR and age (r = 0.41, p = 0.008). For SNR -9 dB there was a significant correlation with age (r = 0.51, p = 0.0007) and threshold (r = -0.37, p = 0.017). Also, with SNR - 12 dB, there was a significant correlation with age (r = 0.54, p = 0.0003). For the more difficult 80% correct criterion, there was a significant correlation with age for -6 dB SNR (r = 0.35, p = 0.026), -9 dB SNR (r = 0.34, p = 0.028), and -12 dB SNR (r = 0.5, p = 0.0009).

To examine at what age declines in performance on the SSIN -12 dB SNR task begin to emerge, a one-way ANOVA revealed a significant effect of age group [F(2,39) = 8.98, p = 0.0006) with post-hoc tests indicating a significant difference between the young and both the middle (p = 0.022) older (p = 0.0004) groups but no difference between the middle and older group (p = 0.29). Similar results were found with -6, -9 and -12 dB SNRs for both the 40% and 80% correct criterion SSIN tasks (not shown for brevity). Thus, performance on the SSIN task was dependent on age, emerging at middle (> 40 years) ages, and not dependent on audiometric thresholds through 4000Hz.

### Temporal and Spectro-Temporal Processing Ability by Age

Shown in the left panels of Figure 6 are scatterplots of individual results for the TFS test for the 30° IPD condition (Part A) and the STM test for each stimulus (Part B). Note that for TFS a smaller (lower) value indicates poorer performance, while for STM a larger (higher) value indicates poorer performance. Solid lines indicate the linear regression trends. The right panels in Figure 6 are box-and-whisker plots of the results for the 3 age groupings: younger adults (n = 13 for both TFS and STM), the middle group (n = 14 for TFS, 17 for STM) and the older adults (n = 15 for TFS, 17 for STM).

**Figure 6.**
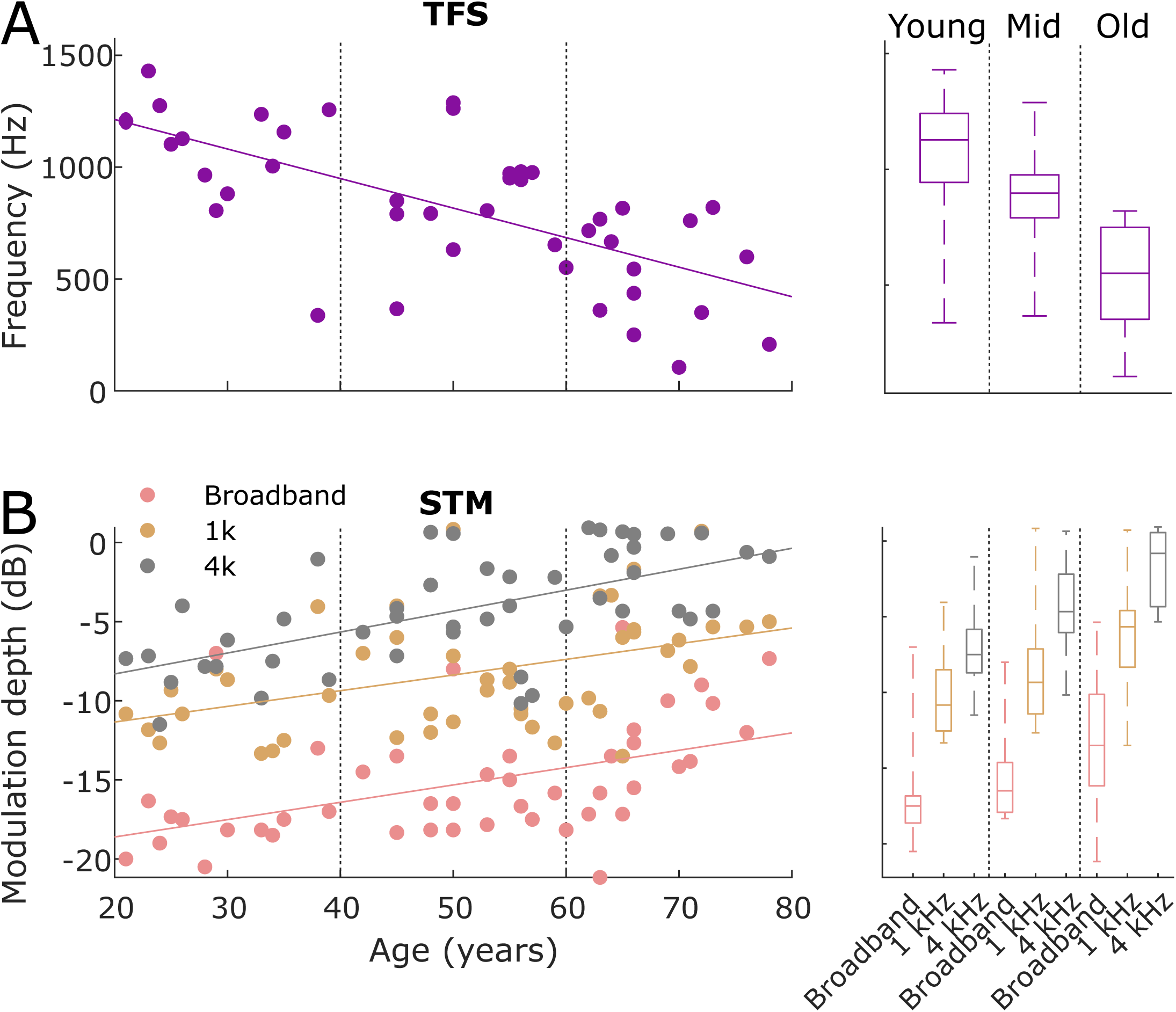
Data for the temporal and spectro-temporal processing tasks. Part A shows results for the TFS test for the 30° IPD condition, and Part B for the STM test for each stimulus type (broadband, 1000 Hz, and 4000 Hz), by age (in years). Left panels show individual data scatterplots, with solid lines indicating the linear regression trends. For TFS (Part A), performance is plotted as the highest frequency at which each subject could discriminate a difference relative to 0°, so a higher number indicates better performance. For STM (Part B). performance is plotted as the minimum modulation depth each listener could detect for each stimulus, so a lower number indicates better performance. Right panels show box-and-whisker plots with the median, lower and upper quartiles, and variability of the data, for 3 age groupings: younger adults (aged 21-39), middle-aged (42-59), and older adults (60-78).

Performance on the TFS task was evaluated at three different IPDs, 30°, 60° and 180°. For TFS with 30° IPD (Figure 6, Part A), although there is variability in individual performance across ages, the linear trend line indicates declining performance with age; similar trends with age were also observed with 60° and 180° IPD with performance improving (i.e., higher threshold frequencies) with increasing IPD (not shown). A two-way ANOVA was conducted that examined the effect of age group and IPD on TFS threshold. There was no statistically significant interaction between the effects of age group and IPD on TFS threshold [F(2, 117) = 0.024, p = 0.99]. Simple main effects analysis showed significant effects of both age (p < 0.0001) and IPD (p < 0.0001). These conclusions result because, while task difficulty increases significantly with decreasing IPD (i.e., smaller threshold frequencies), within each IPD the rate of change of decline in performance with age was the same (i.e., the slopes of decline at each IPD were comparable as a function of age).

Additional partial correlation analyses of TFS with age and hearing threshold at 1000 Hz indicated significant correlation with age, but not threshold, in the 30° (r = -0.66, p < 0.0001), 60° (r = -0.65, p < 0.0001) and 180° IPD (r = -0.57, p < 0.0001) conditions. TFS was also significantly correlated with age, controlling for threshold at 4000 Hz, in the 30° (r = -0.63, p < 0.0001), 60° (r = -0.55, p < 0.0001) and 180° IPD (r = -0.51, p < 0.0001) conditions. To examine at what age group performance on the TFS 30° IPD task begins to decline, a one-way ANOVA revealed a significant effect of age group [F(2,39) = 16.42, p < 0.0001] with post-hoc tests indicating significant differences between the older group and both the young (p < 0.0001) and middle (p = 0.002) age groups, but no difference (p = 0.15) between the young and middle aged. Similar results were found for both 60° and 180° IPD stimuli (not shown). Thus, performance on the TFS task was dependent on age, emerging only in the older age group (> 60 years), and not dependent on audiometric thresholds through 4000 Hz.

For the STM test (Figure 6, Part B), the linear trend lines indicate that the broadband stimulus is the easiest for subjects and 4000 Hz the hardest, and, again, declining performance with age. Partial correlations of STM performance with age and PTA hearing threshold revealed no significant correlation with age or average threshold for the broadband stimulus. Significant correlations were found with age for the 4000 Hz stimulus (r = 0.59, p < 0.0001), controlling for 4000 Hz thresholds, and for 1000 Hz stimulus (r = 0.37, p = 0.012), controlling for 1000 Hz thresholds. To examine at what age performance begins to decline, a one-way ANOVA revealed a significant effect of age on STM 4000 Hz condition [F(2,44) = 15.14, p < 0.0001] with post-hoc tests revealing significant pairwise differences between all 3 age groups (p < 0.05). A similar result was found for the STM 1000 Hz condition (not shown). Thus, performance on the STM task was dependent on age, emerging in middle age (> 40 years), and not dependent on audiometric thresholds through 4000 Hz.

### Correlations Between the Measures

It was of interest to examine whether there was a relationship between results on the spatial acuity/localization and the SSIN tasks for the same subjects, i.e., whether or not the poor performers on one task were also the poor performers on the other task, regardless of age, and how performance on these tasks related to TFS and STM results. To this end, a scatterplot is shown in Figure 7 for individual performances on the most difficult listening condition for each task, because these were where there was greater performance variability shown. Shown are individual MAAs from the localization task for 4000 Hz at -12 dB SNR in the off-axis condition, compared to SSIN scores at -12 dB SNR for each performance criterion (40% and 80% correct). Solid lines indicate the linear regression trends. It can be seen that, although there is fairly substantial performance variability, there is a tendency for poorer performance on one task to correspond to poorer performance on the other task. Pearson correlation analyses revealed low, albeit significant correlations, between SSIN and spatial acuity off-axis with 4000 Hz -12 dB SNR for both the 40% (r = 0.5, p = 0.001) and 80% correct criterion (r = 0.36, p = 0.02). There was also a significant correlation between SSIN scores for the 80% criterion and -12 dB SNR with the off-axis 500 MAA task with -12 dB SNR (r = 0.43, p = 0.004; not shown).

**Figure 7.**
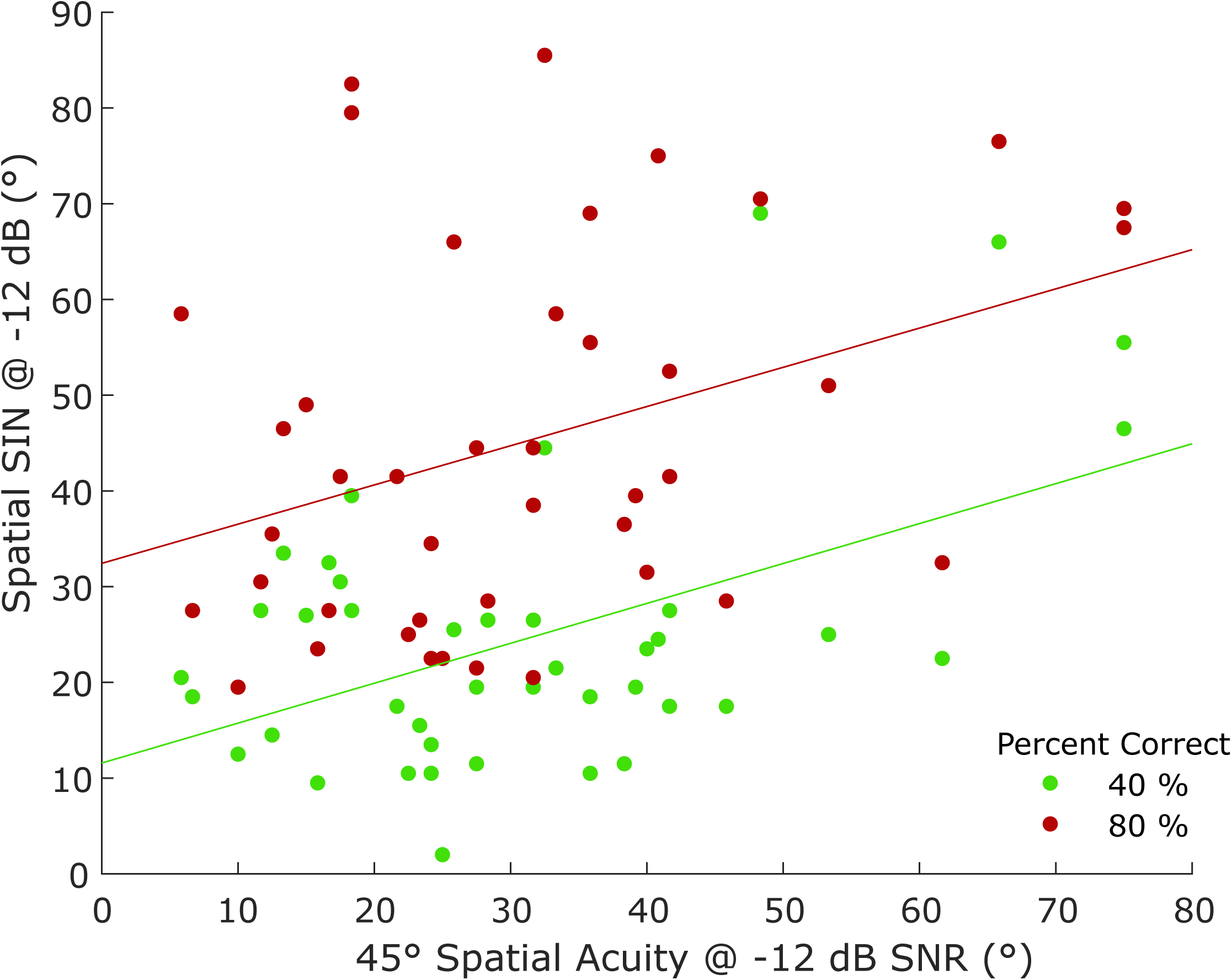
Scatterplot of individual results for spatial-speech-in-noise (SSIN; in degrees separation needed for the speech and noise) for that task’s most difficult SNR (-12 dB) for both the 40% and 80% correct criteria, versus individual results for spatial acuity/localization (minimum audible angle/MAA, in degrees) for that task’s most difficult listening condition (4000 Hz stimulus at -12 dB SNR in the off-axis/45° listening condition).

The top 2 panels of Figure 8 show scatterplots of individual results for the spatial acuity/localization task for the most difficult condition (4000 Hz at -12 dB SNR in the off-axis condition) versus the TFS (Part A) and versus the STM results for each stimulus type (Part B). Solid lines show the linear regression trends. The trend lines illustrate that performance on TFS and STM generally worsens (smaller TFS value and larger STM value) as the MAA for localization worsens. Comparable results are shown in the bottom 2 panels of Figure 8 for SSIN in the most difficult condition (-12 dB SRN for 80% correct).

**Figure 8.**
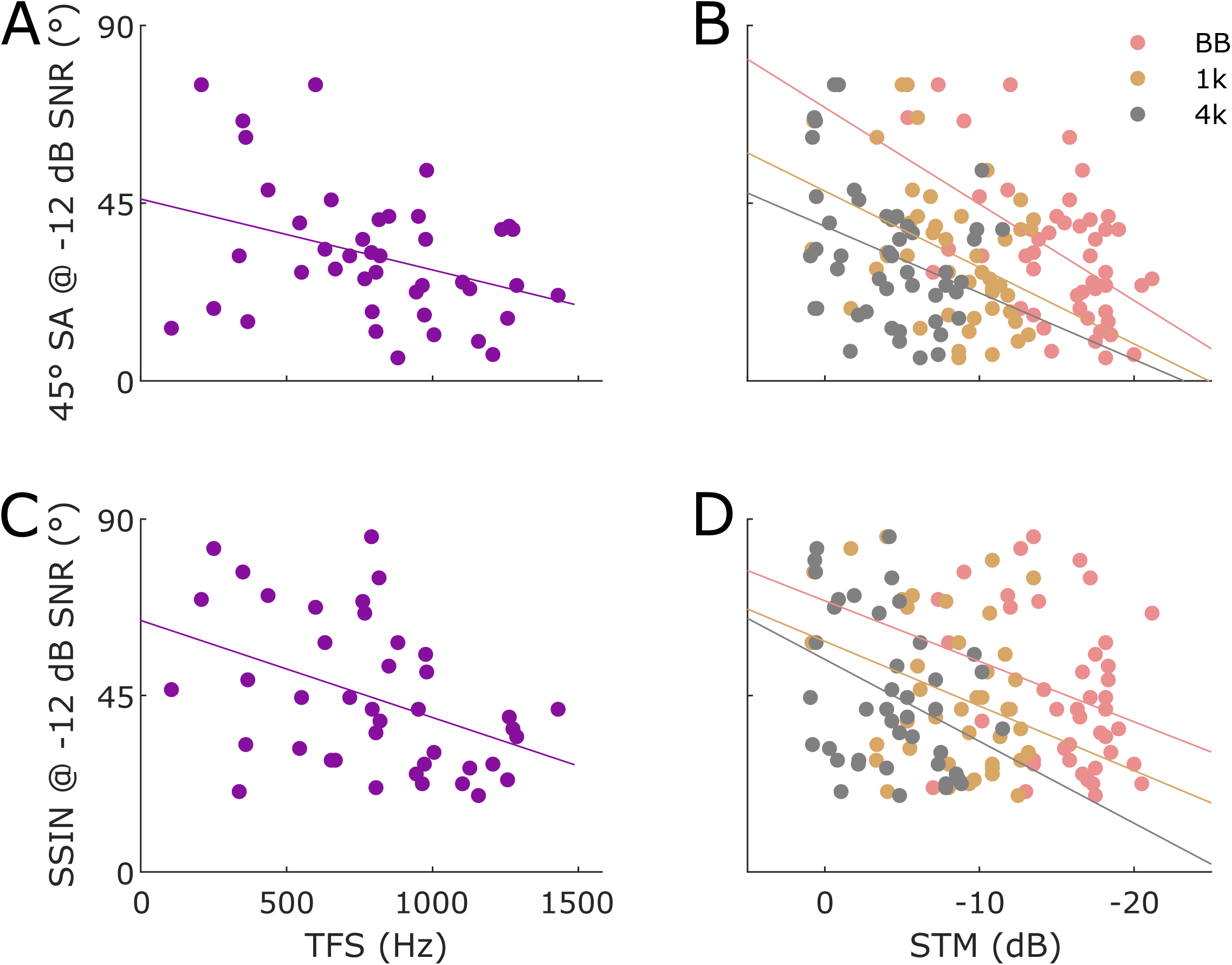
Top 2 panels: scatterplots of individual results for the spatial acuity results (MAA, in degrees) for that task’s most difficult listening condition (4000 Hz stimulus at -12 dB SNR in the off-axis/45° listening condition) versus TFS test results (Part A) and versus STM results for each stimulus type (broadband, 1000 Hz, and 4000 Hz) in the right panel (Part B). Bottom 2 panels: scatterplots of individual results for spatial-speech-in-noise (SSIN; in degrees separation needed for the speech and noise) for that task’s most difficult SNR (-12 dB) for the 80% correct criteria versus the TFS test result (Part C) and versus the STM test for each stimulus type in the right panel (Part D).

Performance on the off-axis MAA with the 4000 Hz stimulus and -12 dB SNR was significantly correlated with performance on TFS, and STM with broadband, 4000 Hz, and 1000 Hz stimuli (r values ranged from 0.34-0.52, p < 0.05). Performance in the SSIN task in the most difficult condition (-12 dB SNR for 80% correct) was significantly correlated with performance on TFS and STM with 4000 Hz and 1000 Hz stimuli (r values ranged from 0.31-0.43, p < 0.05).

Further, we examined correlations between performance and hearing thresholds in the *EHF* range of hearing. Age was significantly correlated with thresholds at all tested EHFs from 9000 to 14000 Hz (r values ranged from 0.76-0.88, p < 0.0001). Performance on tasks including SSIN -12 dB SNR (both 40 and 80% correct), TFS, STM (broadband, 1000 Hz and 4000 Hz) and off-axis MAA for both 500 and 4000 Hz with -12 dB SNR were significantly correlated (r ranged from 0.36-0.57, p < 0.05) with EHFA thresholds. Because EHF thresholds and performance on all tasks were also highly correlated with age, partial correlation analyses were performed with age and average EHF thresholds as factors. Performance on all tasks listed above in this paragraph, *except* the STM tasks, remained significantly correlated with age when controlled for EHF thresholds (r values ranged from 0.31-0.41, p < 0.05). Thus, performance on these tasks was dependent predominantly on age and not on EHF thresholds.

Finally, variables were entered into a multiple linear regression model in order based on the strength of the pairwise Pearson correlations examined above predicting SSIN performance in the -12 dB SNR condition for 40 and 80% correct performance until the r^2^ stopped increasing. The final model included age, TFS for 60° IPD, and MAA for 4000 Hz and -12 dB SNR. The best model predicting SSIN performance at 80% correct was significant [r^2^ = 0.51; F(3, 37) = 12.72; p < 0.0001] with significant prediction by TFS 60° (β = -0.0174; t = 3.11; p = 0.0036) and MAA 4000 Hz -12 dB SNR (β = 0.48; t = 2.05; p = 0.047); age was not significant (β = 0.13; t = 0.94; p = 0.356), even though age helped to increase the r^2^ in the model. Variance inflation factors were < 2.0 indicating multicollinearity was not a factor for this particular model. Results suggest that SSIN recognition scales with the ability of the subjects to use both low-frequency binaural temporal fine structure (60° IPD TFS) as well as higher-frequency binaural envelope cues (4000 Hz MAA).

## DISCUSSION

Overall, the results indicate demonstrable aging effects on spatial hearing in terms of both spatial acuity and spatial speech-in-noise results, primarily in more difficult listening conditions (worse SNRs). Despite being highly correlated with age, hearing thresholds at standard frequencies through 4000 Hz or EHF ranges were not significant predictors of performance on most tasks tested here. For spatial acuity, the conditions in which envelope ITD or ILD cues must be used (4000 Hz, off-axis) resulted in the most notable performance deficits in the older subjects, as expected. A stronger aging effect for an off-axis orientation is consistent with previous research by Briley and Summerfield (2014) who reported declining frontal plane localization ability with aging for the more peripheral regions of frontal space. Our SSIN results are also consistent with previous research that has shown reduced SRM with aging (e.g., Patro et al. 2024; Zobel et al. 2019; Gallun et al., 2013). We did, however, find considerable variability in performances with aging on both spatial hearing tasks, with some older subjects performing as well as young subjects, but others performing much more poorly. This is also consistent with previous research; for example, about half of the subjects over age 60 in Briley and Summerfield (2014) performed about as well as younger subjects but half performed substantially worse.

Finally, a significant relationship was found for performances between SSIN in the most difficult listening condition (-12 dB SNR for 80% correct) and both MAA in the most difficult listening condition (off-axis for the 4000 Hz stimulus at -12 dB SNR), regardless of age, which suggests that both spatial hearing tasks tap into some similar aspect of hearing ability, namely the ability of subjects to use low-frequency fine structure ITDs and higher-frequency envelope ITDs, both of which decline significantly with age.

Following from the above, there was also a clear decline in temporal and spectro-temporal sound processing with advancing age as measured with both the TFS and STM tests, with the strongest relationship between aging and declining performance found using the TFS. When performance on these tasks was compared to performance on the spatial hearing tasks, in the more difficult listening conditions for the latter where variability was greatest, there were some low but significant correlations. These findings suggest that deficits in temporal processing ability may in general underlie poorer performance on spatial hearing tasks regardless of age or hearing thresholds.

In an attempt to reduce the impact of peripheral hearing loss seen with age, we had restricted enrollment to include only up to a mild hearing loss through 4000 Hz, which ensured audibility of stimuli used in the study. Correlations between performance on the tasks and hearing thresholds at 500, 1000, and 4000 Hz reached significance in some cases, although the correlations were generally low, ∼0.3, suggesting that audibility *per se* was likely not a primary factor contributing to the declines in performance on the spatial acuity, SSIN, and TFS/STM tasks with aging. Results of the partial correlation analyses further support this where performance on all tasks tested remained significantly correlated with age even after controlling for standard audiometric range hearing thresholds.

These results depart slightly from the findings of the studies by Bernstein and Trahiotis (2016, 2018, 2019) where they reported that even a slight hearing loss at 4000 Hz of > 7.5 dB resulted in significant performance declines, relative to the subjects with HL < 7.5 dB in tone detection in noise at 500 and 4000 Hz as well as the ability to lateralize 500 and 4000 Hz tones. To be sure, we examined here partial correlations of performance in the off-axis MAA task with 500 and 4000 Hz stimuli at -12 dB SNR (the most difficult condition) with factors of age and hearing thresholds at 4000 Hz. MAAs with both 500 Hz (r = 0.33, p = 0.022) and 4000 Hz (r = 0.5, p = 0.0003) remained significantly correlated with 4000 Hz hearing threshold, controlling for age. Bernstein and Trahiotis used a computational model of peripheral monaural temporal encoding followed by binaural extraction of ITD to reveal that the data could be explained by reduced monaural temporal encoding, e.g., increased jitter, preceding the extraction of ITD. This finding is consistent with more recent computational models of cochlear synaptopathy, where reductions in the number of hair cell to auditory nerve fiber ribbon synapses leads to reduced phase locking in the monaural neural pathways leading to the binaural brainstem nuclei that compute ITD (see also Budak et al., 2022). As discussed above in the Results section, these MAAs were also significantly correlated with EHF hearing thresholds, controlling for age. To the extent to which EHF hearing loss is a proxy for cochlear synaptopathy in more basal low frequency portions of the cochlea (see following discussion), our results here support this hypothesis.

The present results along with the prior studies are consistent with the idea that declines in listening performance on these tasks may be due in part to the sequalae of cochlear synaptopathy (Kujawa and Liberman 2009). Animal studies reveal that both natural aging and even moderate noise exposure in young animals can cause temporary hearing threshold shifts that quickly recover, but permanent loss of auditory nerve to hair cell synapses (Furman et al., 2013; Kujawa and Liberman 2009; Parthasarathy and Kujawa 2018). This aging- and/or noise-induced cochlear synaptopathy *may* be a potential cause for the phenomenon commonly called “hidden hearing loss,” characterized by poor performance on hearing tasks such as speech-in-noise perception despite normal audiological thresholds (e.g., Schaette and McAlpine 2011; Kohrman et al 2019). Indeed, individuals with histories of moderate noise exposure, but normal thresholds, can develop behavioral symptoms consistent with hidden hearing loss (e.g., Füllgrabe et al, 2015). In humans, some loss of synapses begins already in early adulthood (Wu et al., 2019) with additional losses of ∼7.5% per decade (Viana et al., 2015; Makary et al, 2011). In support of this hypothesis, recent animal studies show that noise-induced cochlear synaptopathy causes impaired neural processing of ITD cues in the binaural brainstem (Benson et al., 2025).

However, it is notable that the older subjects in our study did show poorer hearing at 6000 and 8000 Hz on average than younger adults, and also substantially decreased hearing in the EHF range (Figure 1). This is not surprising as it has long been known that EHF hearing loss increases with aging (e.g., see Matthews et al 1997), but it is important to consider that this may be an additional factor underlying the poorer performance by many older subjects on spatial hearing tasks. In fact, correlation analyses on our data showed a surprising number of significant, correlations between average EHF hearing thresholds across 9000-14,000 Hz and performance on the suprathreshold hearing tasks. The strongest correlations were between EHF hearing and the temporal processing tasks, but there were also some significant findings for SSIN and the localization/spatial acuity task. Patro et al. (2024) investigated performance of young to middle-aged subjects (age 20 to age 56) and also reported that EHF-range hearing loss appeared to be a significant contributor to suprathreshold performance deficits in the middle-aged group, producing less SRM on a speech-in-noise test. Further, EHF loss has also been implicated in some limited studies as a potential factor in poor localization ability (Liberman et al. 2016; Brungart & Simpson 2009; Best et al. 2005). For example, Liberman et al. (2016) reported that subjects with EHF hearing loss reported more difficulty localizing the direction of sounds than subjects without EHF loss. Unfortunately, because EHF hearing loss in the current study was seen only in the middle age and older subject groups, it was not possible to determine how much EHF hearing loss contributed to poorer performances on the spatial hearing tasks versus the impact of temporal processing deficits also seen in the older subjects.

Supporting studying EHF hearing loss, Bharadwaj et al. (2019) argued that high-frequency outer hair cell damage in humans almost always occurs concurrently with cochlear synaptopathy in lower-frequency regions. Elevated EHF thresholds could be indicative of cochlear synaptopathy if distortion product otoacoustic emissions are not shown to be affected in lower frequencies. Moreover, EHF hearing loss may be an indicator of the presence of cochlear synaptopathy in more apical cochlear regions subserving lower frequency hearing (Mepani et al., 2021; Mishra et al., 2022). Recently, Mishra et al. (2024) suggested, based on stimulus frequency otoacoustic emissions data obtained at standard audiometric frequencies, that damage in the basal cochlea as revealed by elevated EHF hearing thresholds may be associated with reduced cochlear functioning in more apical regions. Future research is needed in this area to better illuminate this issue.

In conclusion, this study confirms that primary aging effects are seen on average in suprathreshold spatial hearing abilities measured with both a spatial acuity/localization task and a spatial speech recognition in noise task, even when any effect of substantial cognitive deficits is excluded (based on MOCA testing) and/or when hearing loss in the standard audiometric range is excluded (audiometric exclusion criteria) or statistically controlled. The results of this study suggest that these suprathreshold performance deficits may possibly have as an underpinning a decrease in temporal and spectro-temporal processing ability with aging, and also a contribution and/or correlation (i.e., a proxy for cochlear synaptopathy) from EHF-range hearing loss typically seen in older individuals. Notable, however, is that some of the subjects in the oldest age group performed much better than did other older subjects, suggesting an unequal impact of aging across individuals that may be due to any number of factors contributing to degeneration in the auditory system such as the extent of lifetime noise exposure and the presence of cochlear synaptopathy.

In any case, hearing health professionals should remain alert to the fact that some elderly persons may present to the clinic with reports of difficulty understanding speech in noise or localizing the direction of sounds despite having normal or near-normal hearing in the standard audiometric frequency range. Awareness of this problem can be beneficial in counseling and in decisions regarding rehabilitation for these patients. For example, if an active elderly patient complains of suprathreshold hearing deficits but does not show hearing loss of the degree usually considered for hearing aids, very mild gain hearing aids or other assistive hearing devices may still be appropriate for improvement of the SNR that may lead to functional improvement in their daily life. The current data also underscore the importance of measuring EHF hearing thresholds more routinely as part of the audiological evaluation, particularly in older patients.

## Acknowledgment

The authors thank Dr. Melinda Anderson for her contributions to initial study conceptualization and pilot investigations, and Shaylene Denham and Laura Temple for their assistance with administrative tasks. A preprint of this article is available on bioR*x*iv.

## Author Contribution Statement

All authors contributed to this work. D.J.T., C.A.S., K.A.W., N.T.G., and A.K. conceived of and designed the experiments; N.T.G constructed, programmed, and maintained the experimental set-up; K.A.W and C.A.S. performed pilot experimentation, recruited subjects, and collected the data; C.A.S. and D.J.T. performed statistical analyses; N.T.G. drafted figures; C.A.S. wrote the first draft of the manuscript; and K.A.W., D.J.T., N.T.G., an A.K. assisted with critical manuscript revision. All authors discussed the results and implications and commented on the manuscript at all stages.

## No conflicts of interest

This work was funded by NIH-NIDCD R01 DC017924 (MPIs: Tollin and Klug). *The funding organization had no role in the design and conduct of the study; in the collection, analysis, and interpretation of the data; in the decision to submit the article for publication; or in the preparation, review, or approval of the article*.

## List of Abbreviations

ANOVA: Analysis of Variance
dB: decibels
EHF: Extended High Frequency (>8000 Hz)
EHFA: Extended High Frequency Average hearing threshold (9k-14k)
HL: Hearing Level
Hz: Hertz
ILD: Interaural Level Difference
IPD: Interaural Phase Difference
ITD: Interaural Time Difference
LSO: Lateral Superior Olivary
MAA: Minimum Audible Angle
MSO: Medial Superior Olivary
NBN: Narrow Band Noise
PTA: Pure Tone Average hearing threshold (4 frequency 500, 1k, 2k, 4k Hz)
SD: Standard Deviation
SL: Sensation Level
SNR: Signal-to-Noise Ratio
SPL: Sound Pressure Level
SRM: Spatial Release from Masking
SSIN: Spatial Speech-in-Noise
STM: Spectro-Temporal Modulation test
TFS - AF: Temporal Fine Structure – Adaptive Frequency test
2AFC: 2-Alternative Forced-Choice

